# Accelerating Drug Repurposing with AI: The Role of Large Language Models in Hypothesis Validation

**DOI:** 10.1101/2025.06.13.659527

**Authors:** Iratxe Zunzunegui Sanz, Belén Otero-Carrasco, Alejandro Rodríguez-González

## Abstract

Drug repurposing accelerates drug discovery by identifying new therapeutic uses for existing drugs, but validating computational predictions remains a challenge. Large Language Models (LLMs) offer a potential solution by analyzing biomedical literature to assess drug-disease associations. This study evaluates four LLMs (GPT-4o, Claude-3, Gemini-2, and DeepSeek) using ten prompt strategies to validate repurposing hypotheses. The best-performing prompts and models were tested on 30 pathway-based cases and 10 benchmark cases. Results show that structured prompts enhance LLM accuracy, with GPT-4o and DeepSeek emerging as the most reliable models. Benchmark cases achieved significantly higher accuracy, precision, and F1-score (p < 0.001), while recall remained consistent across datasets. These findings highlight LLMs’ potential in drug repurposing validation while emphasizing the need for structured prompts and human oversight.

## I. Introduction

Drug repurposing has emerged as a powerful strategy for accelerating the discovery of effective treatments while significantly reducing the time and costs associated with traditional drug development. Unlike conventional approaches, which require the identification, validation, and clinical testing of entirely new compounds, repurposing leverages existing drugs that have already been approved for other indications. This approach not only shortens the timeline from discovery to clinical application but also mitigates risks related to toxicity and adverse effects, as repurposed drugs have established safety profiles. Moreover, the increasing availability of large-scale biomedical data and computational techniques has further expanded the possibilities for systematic drug repurposing, allowing for the identification of novel drug-disease associations beyond traditional serendipitous discoveries [1].

Traditional drug repurposing approaches often rely on gene-target relationships, where drugs are linked to diseases through shared gene targets. However, this “one-drug-one-target-one-disease” paradigm overlooks the complex biological networks underlying disease mechanisms. To address this limitation, pathway-based drug repurposing has emerged as a complementary and more holistic approach. Biological pathways represent a series of molecular interactions and regulatory mechanisms that govern cellular functions and disease progression. Many diseases share common pathways, meaning that therapeutic interventions targeting one condition may also be effective for another if they influence the same biological processes. This concept forms the basis of pathway-based drug repurposing, which identifies potential treatment candidates by examining disease connections through shared pathways [2].

Despite the promising potential of pathway-based drug repurposing, validating the identified associations remains a critical challenge. Traditional validation methods, such as experimental assays or clinical trials, are resource-intensive and time-consuming. In this context, Large Language Models (LLMs) have emerged as a powerful computational tool to support and enhance the validation of drug repurposing hypotheses.

LLMs, trained on vast amounts of biomedical literature, databases, and clinical reports, can systematically analyze existing knowledge to assess the plausibility of proposed drug-disease associations. These models can extract relevant information from scientific publications, identify supporting evidence, and even generate insights by synthesizing findings from diverse sources. By integrating LLMs into the validation process, researchers can efficiently prioritize the most promising repurposing candidates, guiding subsequent experimental efforts and increasing the overall success rate of drug repositioning [3]. The integration of LLM-based validation with pathway-driven drug repurposing represents a novel and efficient strategy for uncovering new therapeutic applications, bridging computational predictions with real-world biomedical knowledge.

## II. Methods

### A. Overview

This study evaluates the effectiveness of Large Language Models (LLMs) in validating computationally generated drug repurposing hypotheses by systematically testing different prompt engineering strategies. The dataset of drug repurposing cases was generated using a computational method based on analyzing shared biological pathways between diseases and drugs [4, 5]. The detailed prompts and example drug repurposing cases used in this study, as well as the utilized code and obtained results, are available in a publicly accessible GitHub repository [6].

The analysis was conducted in two experimental phases:

- Phase 1 (Prompt Engineering Evaluation): Ten distinct prompt formulations were tested on a set of ten drug repurposing cases (five viable, five non-viable) across four LLMs.
- Phase 2 (Extended Evaluation & Benchmarking): The best-performing prompts and LLMs were further evaluated on thirty additional pathway-based drug repurposing cases. To compare LLM performance on well-established repurposing cases, ten benchmark drug-disease associations (based on historical evidence rather than computational prediction) were included.

To ensure a systematic evaluation, LLM performance was assessed using predefined classification metrics: accuracy, precision, recall, and F1-score. A graphical representation of the methodology is provided in Figure 1.

**Fig. 1.**
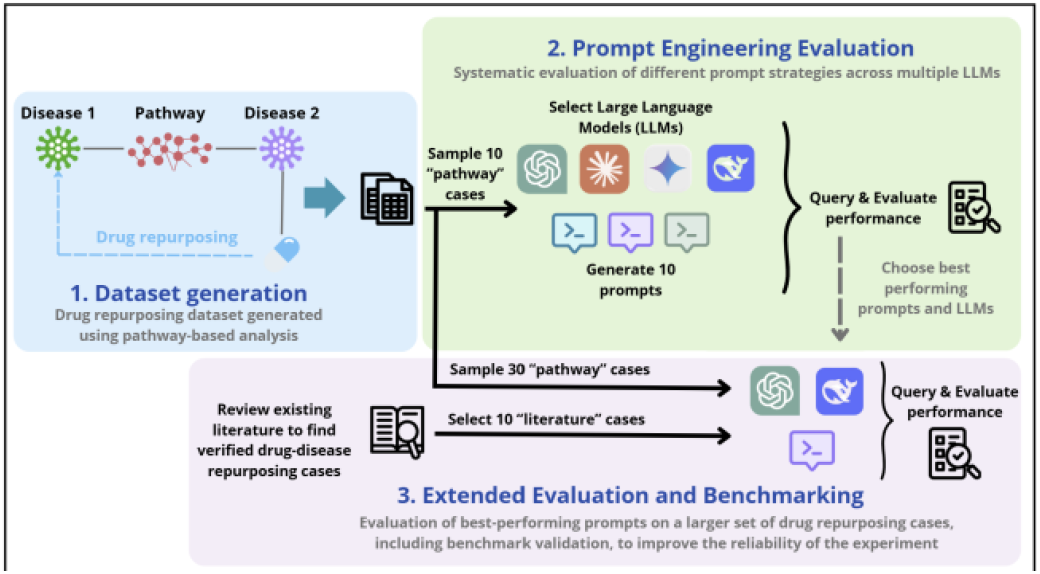
Overview of the methodological pipeline, including dataset generation, prompt engineering, and evaluation process.

### B. Dataset Generation

The identification of drug repurposing cases in which biological pathways played a key role—referred to as DREBIOP (Drug REpurposing based on BIOlogical Pathways)—was carried out through a prior study [4]. Building upon the insights gained from this investigation, a novel drug repurposing approach was developed, leveraging direct relationships between diseases through shared biological pathways [5]. This approach hypothesizes that if Disease 1 and Disease 2 are connected via a common biological pathway, drugs used to treat Disease 2 could be considered as potential candidates for treating Disease 1, based on their shared underlying biological mechanism.

The results obtained from this study led to the identification of a new set of potential drug repurposing cases that required further validation. Specifically, a total of 21,968 disease-drug associations were identified, among which 16,848 represented novel associations between a disease and a potential repurposed drug via a specific biological pathway. At this stage, a new strategy was designed to validate these drug repurposing hypotheses using Large Language Models (LLMs), aiming to enhance the reliability and interpretability of the proposed associations.

To illustrate the types of drug-disease associations included in the dataset, Table I presents a representative sample of pathway-based repurposing cases. Each case includes the original and repurposed indication, the shared biological pathway connecting them, and the viability classification based on literature review. These examples highlight the diversity and biological relevance of the evaluated hypotheses.

**TABLE I.** Sample OF Pathway-Based Drug Repurposing Cases Used IN THIS Study FOR LLM Validation.

| Drug Name | Original Indication | Repurposed Indication | Shared Pathway | Viability |
| --- | --- | --- | --- | --- |
| Ezetimibe | Hyperlipidemia | Familial hypercholesterolemia | WP5109 | Viable |
| Minocycline | Alzheimer's disease | Parkinson's disease | WP5180 | Viable |
| Ephedrine | Obesity | Atherosclerosis | WP5181 | Non-viable |
| Propanolol | Hypertensive disease | Diabetes Mellitus | WP5133 | Non-viable |

The full dataset of 30 pathway-based cases used in this study is available at this study’s GitHub repository (https://github.com/iratxe-zunzunegui/drug-repurposing-validation-LLMs).

### C. LLM selection

To assess the feasibility of LLM-based validation of drug repurposing hypotheses, four state-of-the-art LLMs were selected based on their biomedical capabilities, accessibility, and reasoning efficiency. The chosen models included GPT-4o (OpenAI), Claude-3 (Anthropic), Gemini-2 (Google DeepMind), and DeepSeek (DeepSeek AI).

GPT-4o was selected due to its strong performance in biomedical literature comprehension and structured reasoning, as demonstrated in previous studies evaluating LLMs on medical tasks, including systematic reviews and clinical decision support [7]. Claude-3, known for its robust logical reasoning and contextual depth [8], was included to assess interpretability and consistency. Gemini-2 was chosen for its strong multimodal capabilities, enabling it to process and integrate various data types such as text, images, and audio [9]. Additionally, it leverages Google’s extensive expertise in biomedical research, enhancing its relevance for this study [10]. Lastly, DeepSeek, an emerging open-source alternative, was included to explore whether non-proprietary models can achieve comparable performance in biomedical applications, as recent studies have demonstrated promising results from open-source medical LLMs [11].

Each model was accessed through its respective API or online interface, and their outputs were systematically compared across different experimental conditions.

### D. Evaluation metrics

Performance evaluation was conducted using four standard classification metrics widely employed in machine learning and biomedical NLP tasks [12, 13]: accuracy, precision, recall, and F1-score. These metrics comprehensively assessed model performance in classifying viable and non-viable drug repurposing cases.

For the 30 pathway-based drug repurposing cases, the unweighted versions of these metrics were applied, as the dataset contained an equal distribution of viable and non-viable cases. However, for the 10 benchmark drug repurposing cases, a weighted metric approach was used to account for class imbalance (7 viable, 3 non-viable cases), so that the contribution of each class to the final score was proportional to its frequency in the dataset. This approach ensured that the minority class was not underrepresented in performance calculations.

### E. Prompt Engineering Experiment (Phase 1)

a. *Experimental Design:* To evaluate the impact of prompt formulation on LLM performance, ten distinct prompts were designed and tested on a randomly selected subset of ten drug repurposing cases (five viable, five non-viable) from the dataset described in Section 2.2. Each prompt was applied to all four LLMs, resulting in forty experimental conditions (10 prompts × 4 LLMs). The viability of each drug repurposing case—serving as the ground truth—was determined through a thorough literature search, ensuring that each case was supported or refuted by scientific evidence from peer-reviewed biomedical sources. Evaluation metrics were then applied to quantify model performance.
b. *Prompt Engineering Strategies:* The prompts were designed using a range of natural language processing (NLP) techniques to investigate how different formulations impact response quality and classification accuracy. The key strategies employed included:
  1. *Zero-shot prompting*, where the LLM received a direct query without additional context [14].
  2. *Biological pathway contextualization*, where the model was explicitly informed that the drug-disease association was identified through shared biological pathways.
  3. *Explicit reasoning*, requiring the model to provide logical justification before classifying a case as viable or non-viable.
  4. *Few-shot learning*, where 2 labeled examples were provided before querying the model [14].
  5. *Chain-of-thought prompting*, which guided the model through a structured, step-by-step reasoning process [15].
  6. *Self-critique prompting*, where the model was instructed to assess and refine its response [16].
  7. *Counterfactual reasoning*, which prompted the model to consider potential reasons why a drug might not be viable before classifying it [17].
  8. *Direct literature summarization*, in which the model was tasked with extracting and synthesizing biomedical evidence.
  9. *Expert persona simulation*, instructing the model to assume the role of a biomedical researcher when providing responses [18].
  10. *Hybrid methods*, combining multiple strategies to enhance response quality and reliability.

As an example of how the prompts were applied, consider the case of minocycline, an antibiotic originally used for the treatment of Alzheimer’s disease, which was identified as a potential repurposing candidate for Parkinson’s disease based on shared biological pathways. To evaluate this hypothesis, models were queried using ten distinct prompt formulations. For instance, under the Chain-of-Thought strategy, the model received the following input:

*“This drug repurposing case was identified by analyzing shared biological pathways. Cases generated through this methodology typically involve a high number of pathways, increasing the potential for mechanistic overlap. To determine if minocycline can be repurposed for Parkinson’s disease, follow these steps:*

1. *Identify the primary mechanism of action of minocycline*.
2. *Examine whether this mechanism interacts with key pathological features of Parkinson’s disease*.
3. *Consider existing biomedical literature or clinical trials supporting or refuting this repurposing case*.

*Now, provide your answer beginning with ‘Viable’ or ‘Non-Viable,’ followed by a brief step-by-step explanation and references if available*.*”*

Each prompt encouraged different reasoning approaches and information synthesis. The full set of drug repurposing cases and example prompts is available in the study’s GitHub repository https://github.com/iratxe-zunzunegui/drug-repurposing-validation-LLMs).

### F. Extended Evaluation with Best Prompts and Benchmark Cases (Phase 2)

a. *Experimental Design:* Following the initial prompt evaluation, the four highest-performing prompts were selected based on their F1-score performance. They were then tested on a larger dataset of thirty pathway-based drug repurposing cases (15 viable, 15 non-viable), also randomly sampled from the dataset mentioned in Section 2.2. To determine whether improvements observed in Phase 1 were consistent across a broader dataset, these prompts were applied to the two best-performing LLMs (GPT-4o and DeepSeek).
b. *Benchmark case evaluation:* To further assess LLM performance, a benchmark dataset of ten well-established drug repurposing cases was introduced. These cases were selected based on historical validation in biomedical literature, rather than computational pathway-based predictions, providing an additional reference for model evaluation [1]. The dataset comprised seven viable and three non-viable cases, and weighted evaluation metrics were applied to account for class imbalance. The final comparative analysis assessed LLM performance on the 30 pathway-based cases versus the 10 benchmark cases. Statistical evaluation was conducted to determine the significance of observed performance differences across datasets.

## III. Results and Discussion

### A. Phase 1: Prompt Engineering Evaluation

Table 1 presents the mean evaluation metrics for each LLM (averaged across all prompts) and prompt (averaged across all LLMs), for the 10 drug repurposing cases in Phase 1. The table summarizes accuracy, precision, recall, and F1-score, offering an overview of performance variations across different experimental conditions.

Across all LLMs, GPT-4o demonstrated the highest accuracy (0.83), followed closely by DeepSeek (0.82). Both models consistently outperformed Claude-3 (0.61) and Gemini-2 (0.65), which showed weaker overall classification reliability. Notably, recall was consistently high for all models, exceeding 0.90, indicating that LLMs tend to classify most viable drug repurposing cases correctly. Gemini-2, in particular, achieved the highest recall (1.00), but this was accompanied by a relatively low precision score (0.60), suggesting a higher rate of false positives. Conversely, DeepSeek maintained a more balanced trade-off between precision (0.81) and recall (0.92), while GPT-4o led in overall classification accuracy.

At the prompt level, P4 (Few-shot), P5 (Few-shot with Explicit Reasoning), and P6 (Chain-of-Thought) demonstrated the best accuracy and precision. P6 exhibited one of the highest precision values (0.80) while maintaining perfect recall (1.00), supporting the idea that guiding LLMs through step-by-step reasoning improves classification reliability. Meanwhile, P1 (Zero-shot Prompting) and P3 (Zero-shot Prompting with Explicit Reasoning) performed well in recall but had lower precision, reflecting an increased tendency toward overclassification of viable cases.

A heatmap (Figure 2) visualizes F1-score variation across all LLM-prompt combinations, clearly illustrating how prompt engineering strategies influence model performance. F1 score, which balances precision and recall, was used as the key metric to select the top-performing prompts and LLMs for Phase 2 of the study. As seen in Table 1,-4o (F1 = 0.86) and DeepSeek (F1 = 0.85) emerged as the most reliable models, demonstrating a consistent ability to balance true positives while minimizing misclassification errors. Meanwhile, Claude-3 and Gemini-2 had lower F1-scores (0.71 and 0.74, respectively), confirming their weaker precision in this experiment.

**Fig. 2.**
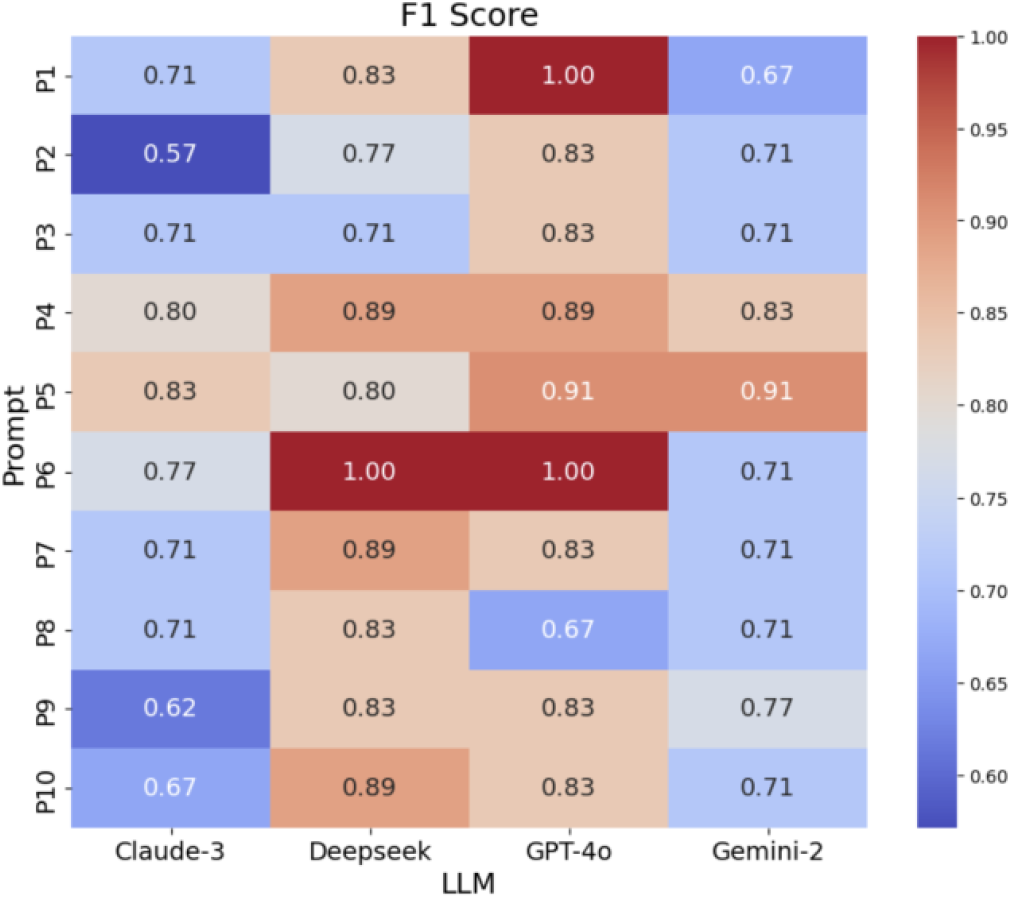
Heatmap of F1 scores for each prompt across the evaluated LLMs. Warmer colors indicate higher performance, with the best-performing prompts and models achieving higher F1 scores.

Among prompts, P6 (F1 = 0.87) was the most effective. P5 and P4 followed closely, achieving F1 scores of 0.86 and 0.85, respectively. These results highlight that providing structured reasoning steps or reference examples improves response quality and model confidence. The effectiveness of Chain-of-Thought and Few-Shot prompting aligns with prior findings in biomedical NLP [19, 20], which emphasize the role of step-by-step reasoning and contextual grounding in improving AI decision-making accuracy.

For this reason, P4, P5, and P6 were selected for further evaluation in Phase 2. Additionally, P1 (Zero-Shot) was also chosen, as it demonstrated relatively strong performance while serving as a baseline comparison to assess the added value of structured prompt formulations. This selection ensured a diverse set of prompt engineering strategies would be tested in the extended evaluation phase.

The results of Phase 1 highlight the importance of prompt engineering in optimizing LLM-based drug repurposing validation. The performance variations observed across models indicate that LLMs require structured prompts to be effective. A key limitation is their tendency to overclassify viable cases, which suggests that human oversight remains necessary, especially when applying LLMs to novel drug repurposing predictions where errors may propagate into real-world decision-making.

### B. Phase 2: Extended Evaluation

Following Phase 1, the four best-performing prompts (P1, P4, P5, and P6) were applied to a larger set of 30 additional pathway-based drug repurposing cases (15 viable, 15 non-viable). This phase aimed to validate the robustness of the selected prompts while narrowing the focus to the two highest-performing LLMs, GPT-4o and DeepSeek. The results, visualized in Figure 3, illustrate the comparative performance of each LLM across the four selected prompts, measured by accuracy, precision, recall, and F1-score.

**Fig. 3.**
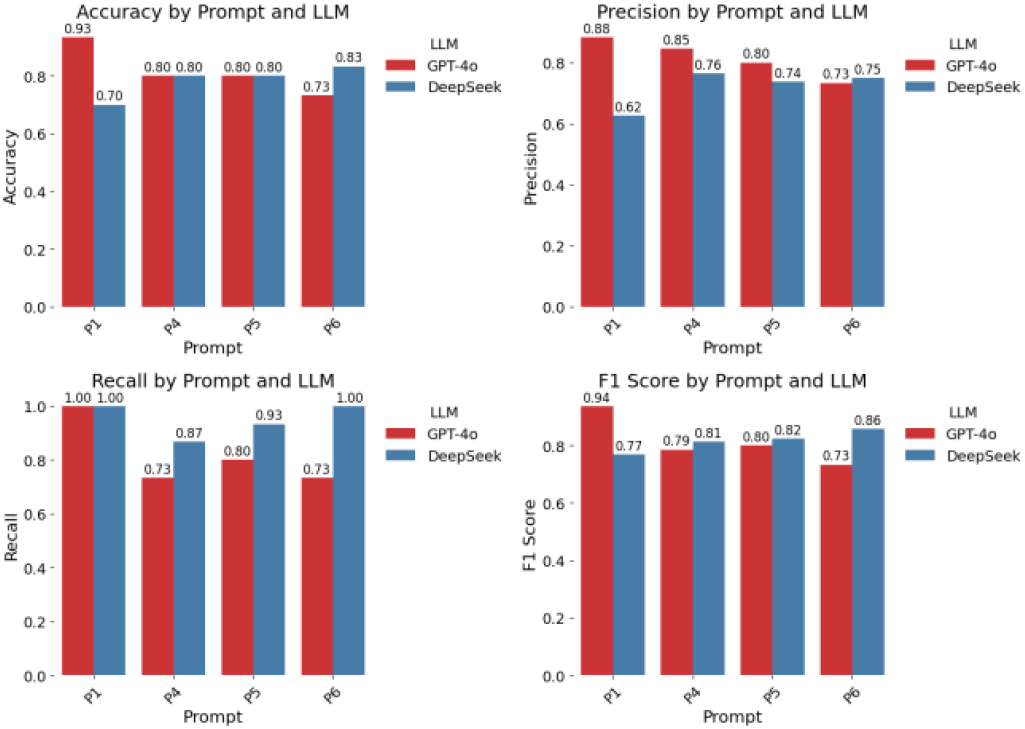
Performance comparison of GPT-4o and DeepSeek on the extended dataset.

GPT-4o maintained the highest accuracy (0.82) and precision (0.82), indicating that it produced more reliable classifications with fewer false positives. In contrast, DeepSeek exhibited slightly lower precision (0.72) but significantly higher recall (0.95), suggesting that it had a greater tendency to classify cases as viable. This behavior reflects a common tradeoff [21] in LLM classification tasks: higher recall improves sensitivity in identifying true positives, but at the risk of increasing false positives. In biomedical applications, such a tendency must be carefully managed to avoid misclassification of ineffective treatments.

Among the prompts, P1 (Zero-Shot) demonstrated the highest recall (1.00), meaning it successfully identified all viable cases. However, its lower precision suggests an increased rate of false positives, which could reduce its reliability in real-world drug validation tasks. P6 (Chain-of-Thought), despite its effectiveness in the previous phase, showed a slight drop in accuracy (0.78). This decline may stem from the complexity introduced by multi-step reasoning, which, while beneficial for interpretability, can sometimes introduce model uncertainty when biomedical information is ambiguous.

The results highlight a key tradeoff between the two LLMs: GPT-4o offers greater predictive reliability for high-confidence drug validation, while DeepSeek’s strong recall may be beneficial in exploratory settings where capturing all potential candidates is prioritized over minimizing false positives. Prompt formulation remains crucial, with structured approaches like P4 (Explicit Reasoning) and P5 (Few-Shot Learning) consistently improving performance. These findings underline the need to carefully balance prompt design and model selection to enhance AI-assisted drug repurposing validation.

### C. Benchmarking Against Established Drug Repurposing Cases

To further assess LLM performance, a benchmark dataset of 10 well-established drug repurposing cases (7 viable, 3 non-viable) was introduced. Unlike the computational pathway-based predictions used in previous phases, these cases were drawn from historically validated drug-disease associations and served as an independent validation set. By comparing LLM performance across both datasets, this phase aimed to determine whether models exhibit greater reliability when classifying well-documented repurpose cases.

The results, presented in Table II, show that both GPT-4o and DeepSeek achieved near-perfect accuracy (0.92) and F1-scores (0.92), demonstrating strong agreement with established biomedical evidence. This high level of precision suggests that LLMs are well-calibrated for cases with strong literature support, reinforcing their potential utility for assisting in systematic reviews and evidence synthesis. However, differences in prompt performance were still observed. P1 (Zero-Shot) and P5 (Few-Shot) achieved the highest F1-scores (0.95), confirming that LLMs excel in well-known cases when presented with minimal or structured context.

**TABLE II.** Mean Performance Metrics Across LLMsand Prompts.

| Model/Prompt | Mean Performance Metrics |  |  |  |
| --- | --- | --- | --- | --- |
|  | Accuracy | Precision | Recall | F1 score |
| Claude-3 | 0.61 | 0.58 | 0.94 | 0.71 |
| DeepSeek | 0.82 | 0.81 | 0.92 | 0.85 |
| GPT-4o | 0.83 | 0.79 | 0.98 | 0.86 |
| Gemini-2 | 0.65 | 0.60 | 1.00 | 0.75 |
| P1: Zero-shot | 0.73 | 0.69 | 1.00 | 0.80 |
| P2: Biological Pathway Contextualization | 0.63 | 0.58 | 0.95 | 0.72 |
| P3: Explicit Reasoning | 0.65 | 0.60 | 1.00 | 0.74 |
| P4: Few-shot | 0.85 | 0.88 | 0.85 | 0.85 |
| P5: Few-shot with Explicit Reasoning | 0.85 | 0.80 | 0.95 | 0.86 |
| P6: Chain-of-Thought | 0.83 | 0.80 | 1.00 | 0.87 |
| P7: Self-critique | 0.73 | 0.71 | 0.95 | 0.79 |
| P8: Counterfactual Reasoning | 0.63 | 0.58 | 1.00 | 0.73 |
| P9: Direct Literature Summarization | 0.70 | 0.64 | 0.95 | 0.76 |
| P10: Persona Simulation | 0.70 | 0.69 | 0.95 | 0.78 |

**TABLE III.** Mean Performance METRICS OF LLMsand Prompts IN Benchmark Cases VS. Pathway Cases AND Statistical Significance OF Differences.

| Metric | Benchmark Mean (LLMs) | Benchmark Mean (Prompts) | Pathway Mean | Statistical Significance |
| --- | --- | --- | --- | --- |
| Accuracy | DeepSeek: 0.925<br>GPT-4o: 0.925 | P1: 0.95<br>P4: 0.90<br>P5: 0.95<br>P6: 0.90 | 0.800 | T-Statistic: 6.35<br>P-Value: 0.0004** |
| Precision | DeepSeek: 0.934<br>GPT-4o: 0.934 | P1: 0.956<br>P4: 0.912<br>P5: 0.956<br>P6: 0.912 | 0.815 | T-Statistic: 6.26<br>P-Value: 0.0004** |
| Recall | DeepSeek: 0.925<br>GPT-4o: 0.925 | P1: 0.95<br>P4: 0.90<br>P5: 0.95<br>P6: 0.90 | 0.875 | T-Statistic: 1.12<br>P-Value: 0.299 |
| F1 score | DeepSeek: 0.920<br>GPT-4o: 0.920 | P1: 0.947<br>P4: 0.893<br>P5: 0.947<br>P6: 0.893 | 0.807 | T-Statistic: 6.27<br>P-Value: 0.0004** |

A notable observation is that precision was higher in benchmark cases (0.93–0.95) compared to pathway-based cases (0.72–0.82), suggesting LLMs show greater confidence in well-established drug-disease associations. Statistical analysis confirmed that accuracy, precision, and F1-score were significantly higher for benchmark cases (p < 0.001). However, recall was similar across both datasets, indicating that LLMs exhibited consistent sensitivity but improved specificity with benchmark cases. This pattern suggests that while LLMs are effective at recognizing positive cases, their confidence in rejecting non-viable cases is higher when dealing with well-documented biomedical associations.

One reason for this may be that benchmark cases align more closely with LLM training data, as they are well-documented, whereas pathway-based predictions introduce more ambiguity. Another key finding is that prompt engineering had a lesser impact in benchmark cases, suggesting that LLMs exhibit greater uncertainty when evaluating novel computational hypotheses. These findings reinforce the importance of human oversight in validating computational hypotheses.

Nonetheless, these findings highlight the promise of LLMs in automating drug repurposing validation and assisting in the identification of novel therapeutic opportunities. The ability of LLMs to systematically assess biomedical literature and provide structured reasoning demonstrates their potential to enhance drug discovery workflows. A compelling example is verapamil, a calcium channel blocker that was newly identified as a potential repurposing candidate for diabetes mellitus through DREBIOP [5]. In this study, the full set of ten distinct prompting strategies was applied to assess the viability of repurposing verapamil.

For example, in the Few-shot with Explicit Reasoning prompt setting, shown in Figure 4, the LLM was given a short instruction followed by two labeled examples (one viable, one non-viable), and finally the target query, which required the LLM to provide a logical justification before deciding whether Verapamil can be repurposed for Diabetes Mellitus. LLMs consistently classified verapamil as a viable repurposing candidate for diabetes throughout various prompt settings—a prediction later verified through manual literature review [22, 23], showcasing the real-world applicability of AI-driven validation.

**Fig. 4.**
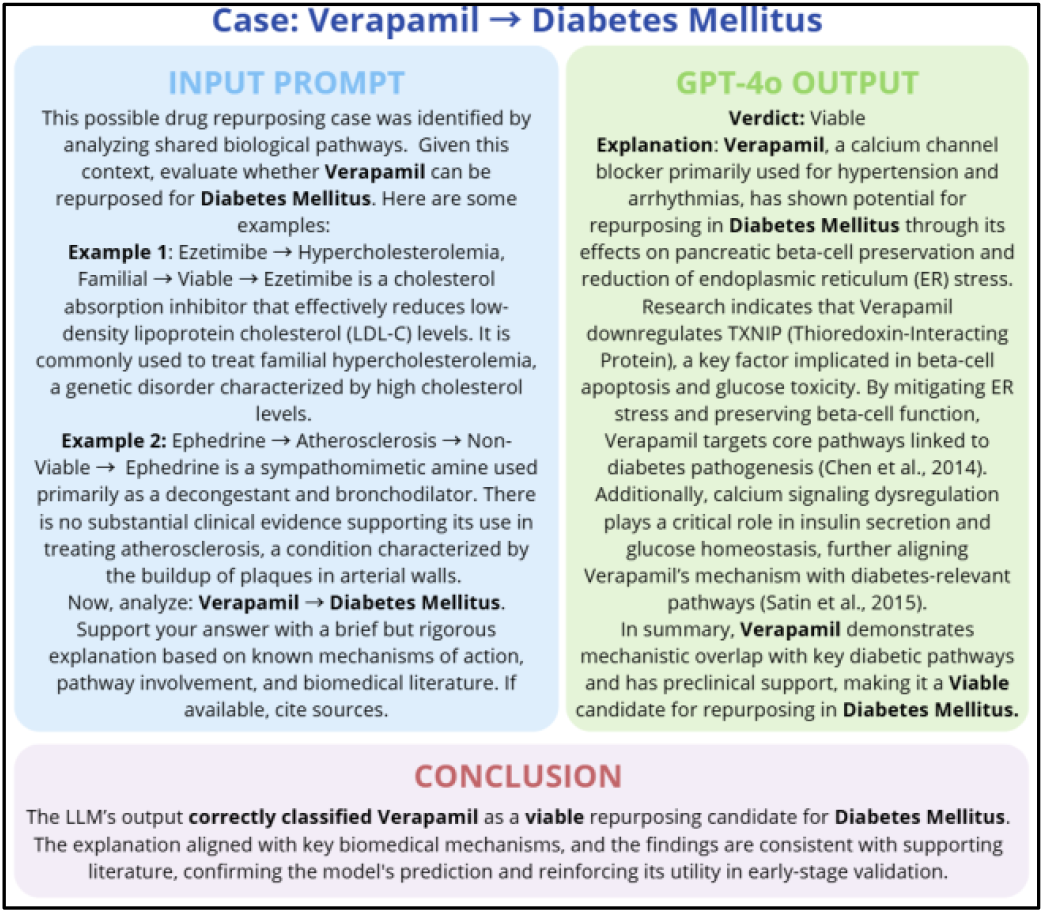
Illustrative example of LLM Evaluation for a viable drug repurposing case

To illustrate a contrasting scenario, Figure 5 includes an example in which the LLM appropriately rejected a non-viable repurposing hypothesis: Paclitaxel, originally indicated for the treatment of atherosclerosis, was evaluated for potential repurposing in obesity, in this case using the Zero-shot prompt strategy. Despite the shared biological pathway, the model concluded it was not a viable candidate, citing its toxicity and absence of supporting literature. This answer was also then reviewed and confirmed manually [24].

**Fig. 5.**
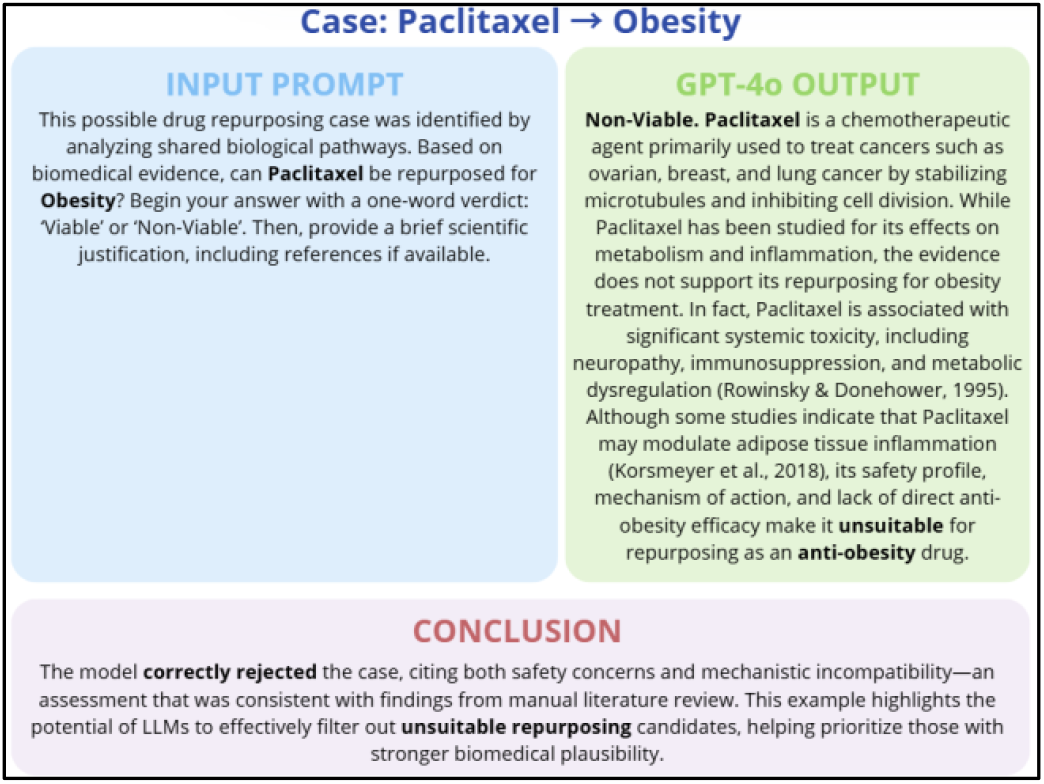
Illustrative example of LLM Evaluation for a non-viable drug repurposing case

These results reinforce the value of integrating LLM-based validation with computational drug repurposing strategies. By filtering out low-potential candidates and highlighting those with promising mechanistic and bibliographic support, LLMs can help speed up the early stages of drug discovery. This enables researchers to focus time and resources on experimentally testing repurposing hypotheses that are more likely to succeed, while deprioritizing those with limited or conflicting evidence.

## IV. Conclusions

This study highlights three major findings. First, prompt formulation significantly impacts LLM performance. Chain-of-thought (P6), few-shot learning (P5), and explicit reasoning (P4) prompts enhanced classification accuracy, whereas zero-shot prompting (P1) demonstrated high sensitivity but also led to increased false positives. Second, GPT-4o and DeepSeek emerged as the most reliable models. GPT-4o achieved the highest precision and maintained balanced performance across all cases, while DeepSeek, although exhibiting slightly lower precision, demonstrated excellent recall, making it a strong open-source alternative. Third, LLMs performed more consistently in benchmark cases than in pathway-based predictions. Greater classification variability was observed in computationally predicted cases, suggesting that LLMs may require additional fine-tuning to generalize more effectively to novel repurposing scenarios.

## V. Future steps and limitations

This study systematically evaluates LLMs and prompt engineering techniques for drug repurposing validation, comparing computational pathway-based predictions with well-established benchmark cases. The findings highlight LLMs’ potential to automate preliminary validation, reducing the time required for manual literature review and allowing researchers to prioritize more promising drug repurposing candidates.

However, several challenges remain. The dataset size is relatively small, requiring expansion to improve statistical generalization and validate findings on a larger scale. Additionally, LLMs were evaluated in their off-the-shelf state, without domain-specific fine-tuning, meaning their performance could improve with biomedical adaptation. Another concern is the tendency of LLMs to hallucinate citations and references, necessitating manual verification to ensure scientific validity. The results also emphasize the importance of structured prompts and human oversight, as inconsistencies in model responses highlight the risks of relying solely on automated outputs.

To enhance the reliability and applicability of LLMs in biomedical research, future work will focus on several key areas. Expanding the dataset will provide a stronger basis for evaluating LLM performance across diverse drug-disease associations. Exploring retrieval-augmented generation (RAG) will allow models to cite real-time biomedical evidence, reducing hallucination issues and improving credibility. Additionally, fine-tuning LLMs on biomedical literature and drug discovery datasets could significantly improve their accuracy, particularly in novel repurposing scenarios, where existing models show greater uncertainty. Addressing these limitations will be essential for optimizing AI-driven drug repurposing validation and ensuring reliable, evidence-based recommendations.

## Acknowledgment

The work is a result of the project “Data-driven drug repositioning applying graph neural networks (3DR-GNN)” that is being developed under grant “PID2021-122659OB-I00” from the Spanish Ministry of Science and Innovation.

